# Mapping Motor Preparation in the Developing Brain: Insights from Contingent Negative Variation and Event-related Mu Rhythm Modulation

**DOI:** 10.1101/2024.03.25.586538

**Authors:** Julia Schmidgen, Theresa Heinen, Kerstin Konrad, Stephan Bender

**Affiliations:** University of Cologne, Univ. Hosp Cologne, Department of Child and Adolescent Psychiatry, Germany; Section Child Neuropsychology, Department of Child and Adolescent Psychiatry, Psychosomatics and Psychotherapy, University Hospital, RWTH Aachen, Germany; JARA-BRAIN Institute II, Molecular Neuroscience and Neuroimaging, Forschungszentrum Jülich GmbH and RWTH Aachen University, Germany

**Keywords:** brain development, motor preparation, contingent negative variation, event-related-potentials, event-related desynchronization

## Abstract

**Introduction:** The motor system shows a pronounced development throughout childhood and adolescence. The analysis of the contingent negative variation (CNV) provides valuable insights into various cognitive and motor processes, underlying cortical sources, and their development across the lifespan.

**Methods:** We investigated the maturation of motor preparation, pre-activation and post-processing in children and adolescents aged 5- to 16- years. EEG Data of 46 healthy right-handed subjects were recorded, using a 64-electrode high density sensor array. Subjects performed a CNV task with a directional warning cue. To assess age related developmental differences of cortical activation, analyses of event-related potentials (ERPs), mu-rhythm (de)synchronization and source analysis were applied.

**Results:** Children showed increased reaction times and committed more errors than adolescent subjects. Motor preparation and post-processing were characterized by a developmental increase of cortical activity related to the supplementary motor area (SMA). Young children showed a pronounced sensory post processing during orienting response (early CNV) that decreased with age. In contrast to previous research in young adults, adolescent subjects showed no contralateral activation of motor areas during motor preparation (late CNV) yet. Furthermore, there was an observed decline in motor post processing with maturation.

**Conclusion:** The results indicate a prolonged maturation of cortical scalp areas associated with motor control up into late adolescence or early adulthood. With age, the activation of mid-frontocentral regions associated with the SMA becomes more pronounced during motor planning and response evaluation. Qualitatively distinct cortical activation patterns of young subjects suggest immature supplementary-, pre- and primary motor areas and might be a primary cause for age-related increasing efficiency of motor action control.

## 1 Introduction

Cortical development in childhood and adolescence underlies complex restructuring processes characterized by extensive anatomical (Huttenlocher, 1979; Shaw et al., 2008) as well as functional changes of the cerebral cortex (Thatcher, 1992). Gaining proficient motor skills stands as a key cornerstone in human development; however, there is limited understanding regarding the relationship between brain development and the acquisition of motor abilities.

The motor system engages in various processes when preparing for movement. This includes the selection of the involved muscles, determining the required contraction force, and arranging the temporal sequence of movements. Changes in brain activity prior to a movement reflect neural processes associated with movement preparation and execution (Deecke et al., 1969). Thereby, it was shown that the supplementary motor area (SMA) plays a crucial role in planning and coordinating voluntary movements (Roland et al., 1980; Tanji, 2001) and that SMA activity has a modulatory effect on the output of the primary motor cortex (M1) (Côté et al., 2020).

Maturation of functional activity related to motor network pre-activation can be studied by analyzing age-dependent changes of event-related-potentials (ERPs) prior to the execution of a movement. The negative potential arising between a warning stimulus (S1) and a behaviorally relevant imperative stimulus (S2) is described as contingent negative variation (CNV) and is associated with processes of movement preparation and attention allocation (Rockstroh, 1982; Walter et al., 1964). The imperative stimulus follows the warning stimulus in fixed time intervals and requires a fast motor response. To efficiently process the imperative stimulus, the neuronal network responsible for initiating the chosen response movement must be pre-activated. Since the CNV is based on a predictable, external cue, it allows to temporally disentangle early response selection and preparation processes, i.e., the recruitment and selection processes of the specific neuronal networks required for a fast response (Gomez et al., 2003).

The CNV consist of an early low, negative component that typically follows the warning stimulus with a latency of 550 to 750 ms and a more pronounced negative late component that precedes the behaviorally relevant imperative stimulus. The early CNV (abbreviated as iCNV; initial CNV; “O-wave”) is suggested to reflect an orienting response (Rockstroh, 1982) and to originate from the SMA and the anterior cingulate cortex (ACC) (Cui et al., 2000; Gomez et al., 2001). The late CNV (abbreviated as lCNV; E-wave) is associated with preparatory processes and stimulus anticipation (Rohrbaugh et al., 1976) observed as pre-activation of the contralateral motor areas. Furthermore, the CNV contains a post-movement potential known as post-imperative negative variation (PINV). This slow negative potential can be studied to evaluate processes related to motor performance evaluation (Klein et al., 1996).

The CNV has been extensively analyzed to study pathophysiological differences between control subjects and subjects with various neurological and mental disorders. However, healthy maturation related to preparatory brain activity remains largely unexplored in terms of its underlying developmental mechanisms. Electrophysiological studies comparing late CNV potentials of children and adolescents showed increasing negativity over pre- and primary motor areas (Bender et al., 2005; Bender et al., 2002; Gomez et al., 2003). The results were interpreted as immaturity of the frontal cortex. However, it remains unclear if absent or low pre-movement negativity in young children directly reflects activity states of the underlying cortical areas and to what extent the supplementary motor area contributes to early response selection, especially in young children.

EEG research of the early CNV component revealed more contradictory results. It was shown that the early CNV exhibits modality-specific characteristics. Specifically, it shows higher amplitudes in response to auditory warning stimuli (S1) compared to visual warning stimuli. Concerning the maturation of early CNV, studies focusing on auditory CNV paradigms have reported decreasing amplitudes over frontal electrodes during childhood and adolescence (Bender et al., 2005), whereas investigations involving visual CNV paradigms have indicated rising frontal early CNV amplitudes within the same age range (Jonkman et al., 2003). These findings suggest that S1 post-processing exhibits a modality specific contribution to the early CNV, showing stimulus dependent developmental differences. Simultaneously, an S1-modality independent component results in increasing fronto-central amplitudes during response selection processes (Jonkman, 2006; Jonkman et al., 2003).

Besides electrophysiological ERP data, changes of brain oscillatory activity of different frequency bands can be analyzed to study task related power changes. Changes within the alpha band are typically observed in motor-related brain areas during processes of motor planning (Pfurtscheller & Lopes da Silva, 1999).The analysis of alpha power enables the modulation of more widespread cortical dynamics and provides complementary information to the analysis of evoked potentials (Claudio Babiloni et al., 1999). A relative increase of frequencies related to the alpha rhythm, commonly assessed in a range of 8 to 12 Hz, refers to alpha band event-related-synchronization (ERS) and is suggested to reflect active processes of inhibition (Jensen & Mazaheri, 2010; Klimesch et al., 2007). Alpha band event-related-desynchronization (ERD) is related to a decrease of oscillatory activity and an increased activity of underlying brain areas (Niedermeyer, 1997). Even though analysis of the alpha rhythm provides additional insights into maturational processes, CNV induced oscillatory changes have been studied rarely in the context of healthy brain development.

The primary aim of this study was to investigate maturational changes of attention-allocation, movement preparation and motor program pre-activation. We collected EEG data and movement performances of a sample consisting of 46 children and adolescent subjects aged 5- to 16-years. Using a visual CNV paradigm, we analyzed cortical activation prior to a predictable, externally triggered movement. To maximize response selection processes, we chose a directional warning stimulus S1 (arrow pointing to right or left side) that indicated the movement side required for the motor response to the imperative stimulus S2. ERP data analysis, alpha-band desynchronization examination, and source localization were conducted to explore developmental processes at various levels.

## 2 Material and Methods

### 2.1 Subjects

A total of 46 typically developing subjects in the age range of 5- to 16-years were recruited. For details regarding demographic characteristics, see Table 1. All participants had no history of motor impairments or any neuropsychiatric condition. A diagnostic interview was used for all subjects to assess and exclude neuropsychiatric disorders (Kinder-DIPS; Adornetto et al. (2008)). The study included only right-handed participants (Edinburgh Handedness Inventory, Oldfield (1971)) that had an IQ score of at least 70 (WISC V; Wechsler, D. (2017)). Participants with an individual or a family history of epilepsy, severe or acute psychiatric diseases, neurological or non-correctable visual impairments were excluded from the study. Subjects were not permitted to take any psychoactive or antipsychotic drugs affecting the central nervous system. Two participants had to be excluded from the analysis due to high artifact levels.

**Table 1.** Sample characteristics.

| Age Group (Years) | N | Mean age (± SD) |
| --- | --- | --- |
| 5-8 | 12 (5 m, 7 f) | 7.27 (± 1.14) |
| 9-12 | 17 (7 m, 10 f) | 10.68 (± 1.34) |
| 13-16 | 15 (7m, 8 f) | 14.72 (± 1.13) |

The experiments were performed in accordance with the Declaration of Helsinki. In an age-adjusted information letter as well as in a personal briefing, participants and parents were informed about the procedure of the measurements and the possibility to terminate participation at any time without giving reasons. Before participation, participants and their parents signed written informed consent.

### 2.2 Experimental procedure

The software package *Presentation* (Version 10.3, Neurobehavioral Systems Inc., Albany, CA) was used to generate a task-related program with alternating visual stimuli displayed on a monitor at a distance of 90 cm from subjects. To prevent any distractions, the light was dimmed, and the noise level was reduced to the highest possible minimum. To minimize distracting eye movements, a fixation cross was displayed in-between visual stimuli. Subjects sat in a comfortable position on a skid-proof chair, reducing muscle activity as much as possible to prevent interfering contractions.

### 2.3 Behavioral CNV task paradigm

Subjects performed a visual CNV (Fig. 1) task with 50 trials, respectively for each response side. The warning stimulus (S1) was presented as a black arrow on a white background pointing to the left or right side. The imperative stimulus (S2) was presented as a colored sheriff on a white background. Both stimuli were displayed for 150 ms, interstimulus intervals were set to 3.05 seconds, pseudorandomized intertrial intervals varied from 3 to 6 seconds.

**Fig. 1.**
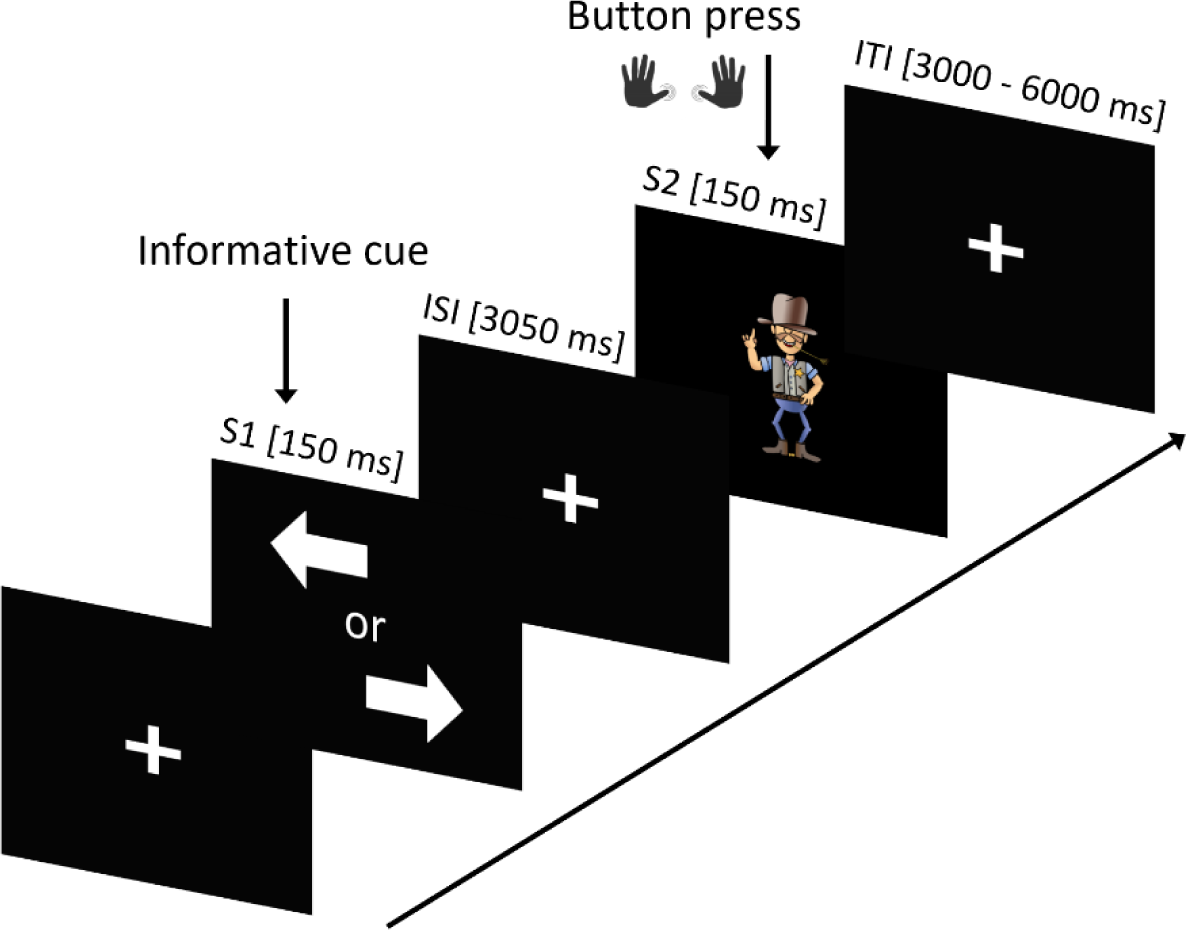
Contingent negative variation (CNV) task paradigm with a directional warning stimulus S1 and a behaviourally relevant imperative stimulus S2.

The warning stimulus indicated the side of the required button press, whereby arrows were presented in a pseudorandomized order. Subjects were instructed to respond to the imperative stimulus S2 as fast as possible by pressing either the right or left button on a German standard keyboard (ctrl for left button press, enter on the numeric keypad for right button press) with the left or right thumb.

### 2.4 EEG

EEG was recorded using a 64-channel BrainAmp system (BrainProducts, Munich, Germany) and Brain Vision Recorder software (BrainProducts). Elastic EEG caps with direct current sintered Ag/AgCl disc electrodes (BrainProducts) were selected based on head sizes. Electrodes were named based on their location on the scalp, consistent with the international 10-20 system, electrode impedances were kept below 5 kΩ. Additionally, EOG electrodes were positioned under the left and right eye and on the nasion. The sampling rate was set to 5000 Hz, electrode CZ was used as recording reference.

### 2.5 Electromyography

Surface EMG was recorded using self-adhesive silver-silverchloride electrodes in a belly tendon montage respectively for the left and right hand. To record thumb movement, active electrodes were placed on the adductor pollicis muscle, reference electrodes were attached to the exterior proximal phalanx of the thumb. The ground electrode was placed on the inner forearm.

### 2.6 Signal Preprocessing

EEG and EMG data were preprocessed using the BrainVision Analyzer2 software (BrainProducts, Munich, Germany). To reduce large file sizes, data were downsampled to 500 Hz. EEG data were re-referenced to an average reference. For the analysis of CNV characteristics, EEG data were segmented into epochs of 7.5 seconds (from 800 ms before warning stimulus to 3000 ms after imperative stimulus), respectively for left and right button press condition. Only trials with correct responses within the time window of 100 to 1500 ms following S2 were included in further analysis. Muscle artifacts were rejected by visual inspection, independent component analysis (ICA) was used to remove artifacts evoked by eye movement (Mennes et al., 2010). Baseline correction was set from 500 to 0 ms preceding the warning stimulus (S1), and signals were filtered digitally (50 Hz notch filter). In order to prevent slow drift effects from effecting EEG data, linear DC detrend was applied. DC detrending had no systematic effect on EEG data. Averages were calculated, respectively for each experimental condition.

### 2.7 Data analysis

#### 2.7.1 Behavioral data

Reaction times were calculated as mean time between the presentation of S2 and the button press of correct response trials. Error rates were calculated from sum of no or false alarms (button press between S1 and S2) and incorrect button presses in response to S2 (button press on wrong movement side).

#### 2.7.2 Event-related potentials (ERP)

Topographical distribution and CNV waveforms of evoked activity were analyzed for the different CNV components, respectively for left and right response condition. iCNV was defined as the mean value of a 200 ms time window around the maximum amplitude at mid-frontocentral electrodes (Fz, FCz’, FC1’, FC2’) between 550 and 1400 ms following S1 (Bender et al., 2002; Böcker et al., 1990; Kropp et al., 1999). Data inspection and iCNV latency analysis revealed a pronounced, continuous latency shift of the early component, especially in younger subjects (see Figure 3A). Consequently, the iCNV peak detection time window was extended from 550 - 750 to 550 - 1400 ms after S1 to prevent a shadowing of maturational differences in iCNV amplitudes. The lCNV component was calculated as the mean voltage of the interval of 200 ms preceding the imperative stimulus S2 (Böcker et al., 1990). The PINV was defined as mean value of a 200 ms time window around the maximum amplitude between 500 and 1500 ms following the imperative stimulus (S2). Electrodes Cz, FCz′, FC1′, FC2′ were used to analyze midfronto-central activity (Cui et al., 2000; Gerloff et al., 1998), electrodes C4, CP4′ CP6′, respectively C3,CP3′, CP5′ were used to analyze motor activity over central areas related to the right and left primary motor cortex (Gerloff et al., 1998).

#### 2.7.3 Alpha event-related desynchronization (alpha-ERD)

We analyzed alpha band desynchronization to study movement related changes of cortical oscillations in the frequency range of 8 to 12 Hz. EEG studies in infants have indicated that lower frequency ranges (6-8 Hz, theta rhythm) can be considered comparable to the mu-rhythm observed in adults (Cochin et al., 2001). However, there is no clear evidence that theta rhythm in infants and mu-rhythm in adults represent equal processes. Furthermore, Berchicci et al. (2011) observed that a shift of “central alpha” activity was most pronounced in the first years of life and reached a frequency of 9 Hz at the age of 4 years. Since our subject sample started at the age of 5 years and to ensure comparability between young children and adolescent subjects, we analyzed common alpha band ERD of 8 to 12 Hz.

Alpha ERD was calculated based on reference free (current source density) EEG data. The same time windows and trials as in ERP analysis were used for analysis of alpha ERD. The signal was bandpass filtered in the frequency range of 8 to 12 Hz, subsequently squared and averaged across segments (Pfurtscheller & Da Silva, 1999). ERD output data was normalized, i.e., baseline corrected and rescaled relative to mean value within the reference interval of 1000 ms prior to the onset of the imperative stimulus S1. ERD magnitude at each electrode was expressed as percentage changes of the instantaneous power.

#### 2.7.4 Analysis of lateralization (event-related potentials, alpha ERD)

To analyze lateralized activity, the lateralized portion of event-related potentials and alpha ERD was computed, taking into account both right and left button press trials for the same time windows as in ERP and alpha ERD analysis.

The calculation is known from the double subtraction method of the lateralized readiness potential (LRP, see de Jong et al. (1988)). The method involves subtracting the average ERP/alpha ERD of ipsilateral electrodes (e.g., left centro-parietal area for left response condition and right centro-parietal area for right response condition) from the average ERP of contralateral electrodes (e.g., right centro-parietal area for left response condition and left centro-parietal area for right response condition). Subsequently, the mean of the differences obtained for both button press sides was calculated.

#### 2.7.5 Current source density (CSD) analysis

To enhance spatial resolution (Nunez et al., 1994) and identify locations of current sources involved in iCNV generation, we used CSD analysis. For CSD estimation, spherical spline interpolation method was applied before surface Laplacian based on the EEG voltage distribution was used. CSD data is indicated as µV/m^2^. Topographical distribution of cortical activity over mid-frontocentral areas (Cz, FCz′, FC1′, FC2′), which is likely to originate from the SMA (Pfurtscheller et al., 2003), was analyzed.

#### 2.7.6 Current Standardized shrinking LORETA-FOCUSS (SSLOFO)

The inverse SSLOFO (standardized shrinking LORETA-FOCUSS) algorithm was used to precisely reconstruct cortical sources (Liu et al., 2005). The algorithm combines multiple techniques to improve source localization abilities. Initially, a low resolution sLORETA image is calculated. To improve spatial resolution, the re-weighted minimum norm of FOCUSS is applied. “Standardization” technique is used to enhance the ability of localization, as in sLORETA. Further insights into the specifics of the SSLOFO algorithm, along with comprehensive performance comparisons against other prevalent algorithms, can be found in Liu et al. (2005).

#### 2.7.7 Statistical analysis

Statistical analysis of the data was performed separately for each CNV component, after confirming the well-known interaction between CNV component and scalp area, which indicates different topographies for the CNV components. To examine maturational changes, data were divided into age groups (5- to 8-year-olds (n = 12), 9- to 12-year-olds (n =17) and 13- to 16-year-olds (n =15)). This approach enhances the capabilities to investigate interactions between age-dependent development and other factors, such as scalp area. Repeated measurement analysis of variance (ANOVA) with the within-subject factors movement side (right, left) and scalp area (mid-frontocentral, right and left centro-parietal) and the between-subject factors age group (5- to 8-year-olds, 9- to 12-year-olds and 13- to 16- year-olds) and gender (male, female) were applied to investigate ERP latencies for iCNV and PINV as well as ERP and alpha-ERD amplitudes for iCNV, lCNV and PINV. Pairwise comparisons of estimated marginal means, including Šídák correction for multiple testing, were integrated in the linear mixed model to test for significant differences across the different age groups and scalp areas. For ANOVA testing and pairwise comparisons of estimated marginal mean, values of *p* < .05 were considered statistically significant.

Multiple t-tests were used to identify areas with activity that significantly differed from the baseline. To address the issue of multiple comparisons, an alpha correction (Bonferroni Correction) related to the tested scalp regions was applied. Due to variations in the number of multiple comparisons across observed parameters, corresponding statistically significant p-values are reported directly in relation to each parameter.

#### 2.7.8 Regression analysis

To investigate developmental effects on the dependent variables, regression analyses with the predictor age were performed. Due to developmental trajectories showing curvilinear age effects on certain dependent variables (i.e., more pronounced changes in younger children) (Fietzek et al., 2000; Klein, 2001), regression analysis was performed either as linear regression using age as predictor (*y* = *a* + *b* * age; *y* = predicted original data; *a* = constant; *b* = regression coefficient) or as non-linear regression using age-1 as predictor (regression equation *y* = *a* + *b* * age^−1^). An exploratory analysis of the scatterplots was used to determine if data showed a linear relationship between age and a dependent variable, i.e., a constant development of the variable throughout the examined age range of 5 to 16 years, or a non-linear influence of the factor age on the dependent variables, i.e., more pronounced development in younger (or older) subjects. The most suitable regression model (predictors age or age^-1^) was chosen based on data examination respectively for each variable.

## 3 Results

### 3.1 Behavioural task performance (reaction time)

Behavioral task performance across age groups was investigated by analyzing error rates and reaction times. The ANOVA analysis on the dependent variable total response error rate, with the between-subject factors of age group and gender revealed significant differences between the age groups (F(2, 38) = 5.75, *p* = .007). Post-hoc Tukey-HSD comparisons indicated that the oldest age group of 13- to 16-year-olds (3.93 ± 3.33) performed significantly better than the youngest age group of 5- to 8-year-olds (13.75 ± 11.01, *p* = .009). The middle age group of 9- to 12-year-olds (10.76 ± 8.25, *p* = .056) exhibited only a trend toward increased errors compared to the oldest age group and no significant differences compared to the youngest age group. The most frequent error observed was a button press during the intertrial interval, occurring between the warning stimulus S1 and the imperative stimulus S2. Only trials with a correct response to S2 were included in further analysis.

Regarding reaction times, ANOVA analysis with age group and gender as between-subject factors showed a trend toward differences among the age groups (F(2, 38) = 2.96, *p* = .064). Tukey-HSD post-hoc comparisons indicated that the youngest age group (443.02 ± 117.42 ms) showed slower reactions compared to the middle (342.03 ± 112.91 ms, *p* = .046) and a trend compared to the oldest age group (369.83 ± 114.12 ms, *p* = .05). These results suggest that younger subjects exhibited more impulsive and slower response reactions compared to older subjects.

### 3.2 Early orienting response and motor preparation (initial contingent negative variation)

#### 3.2.1 Latency of iCNV component

The initial component of the CNV potentials was used to investigate developmental changes during childhood and adolescence related to early movement preparatory processes.

The repeated measures ANOVA revealed a significant effect of the factor age on iCNV latencies (F(2, 38) = 4.84, *p* = .013). The delayed onset of early mid-frontocentral negativity in younger subjects is illustrated in Figure 2A, 2B and 3A. For mean latencies with standard deviations, see supplementary material, Table 2.

**Fig. 2.**
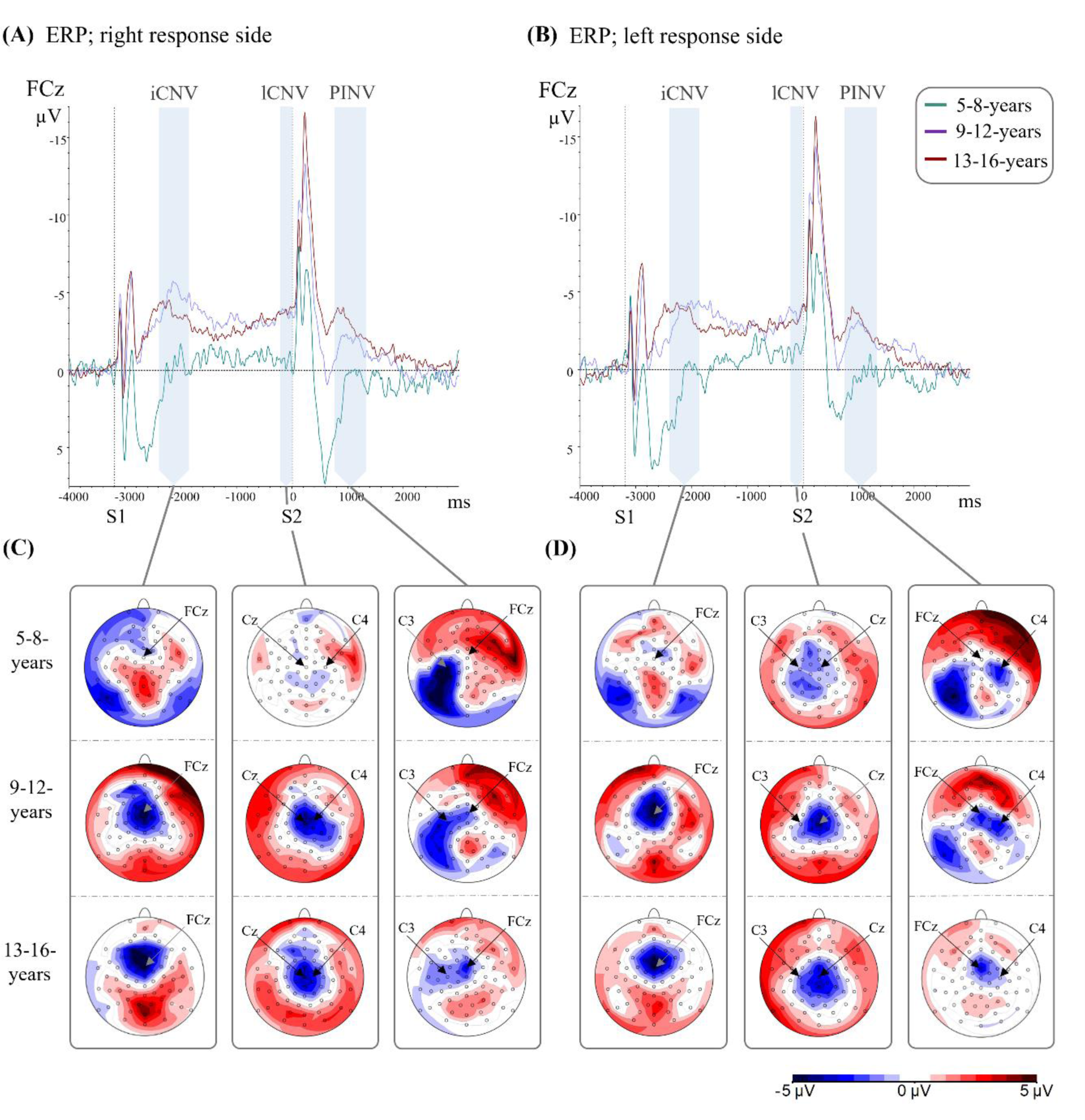
Grand average CNV waveforms and topographic voltage maps, separately displayed for each age group to demonstrate maturational differences. **(A)** CNV course of the right and **(B)** left response condition at FCz (right), respectively for 5- to 8-year-old subjects (green line), 9- to 13-year-old subjects (purple line) and 13- to 16-year-old subjects (red line). The first vertical dashed line indicates the presentation of the warning stimulus S1 (directional arrow), the second vertical dashed line indicates the presentation of the imperative stimulus S2. Time windows for iCNV, lCNV and PINV are highlighted in light blue. Please note that voltage scales are presented upside down with negative values going upward. See for more detailed amplitude values supplementary material, Table 1. **(C)** Voltage maps of each CNV component for the right and **(D)** left response condition, displayed as average of 5- to 8-year-old (top), 9- to 12-year-old (middle) and 13- to 16-year-old subjects (bottom). Maps are scaled from -5 µV (blue) to +5 µV (red). Older subjects showed increasing negativity of mid-frontocentral areas in each component. Most pronounced differences were observed between the age group of 5- to 8-year-old and 9-to 12-year-old subjects.

**Fig. 3.**
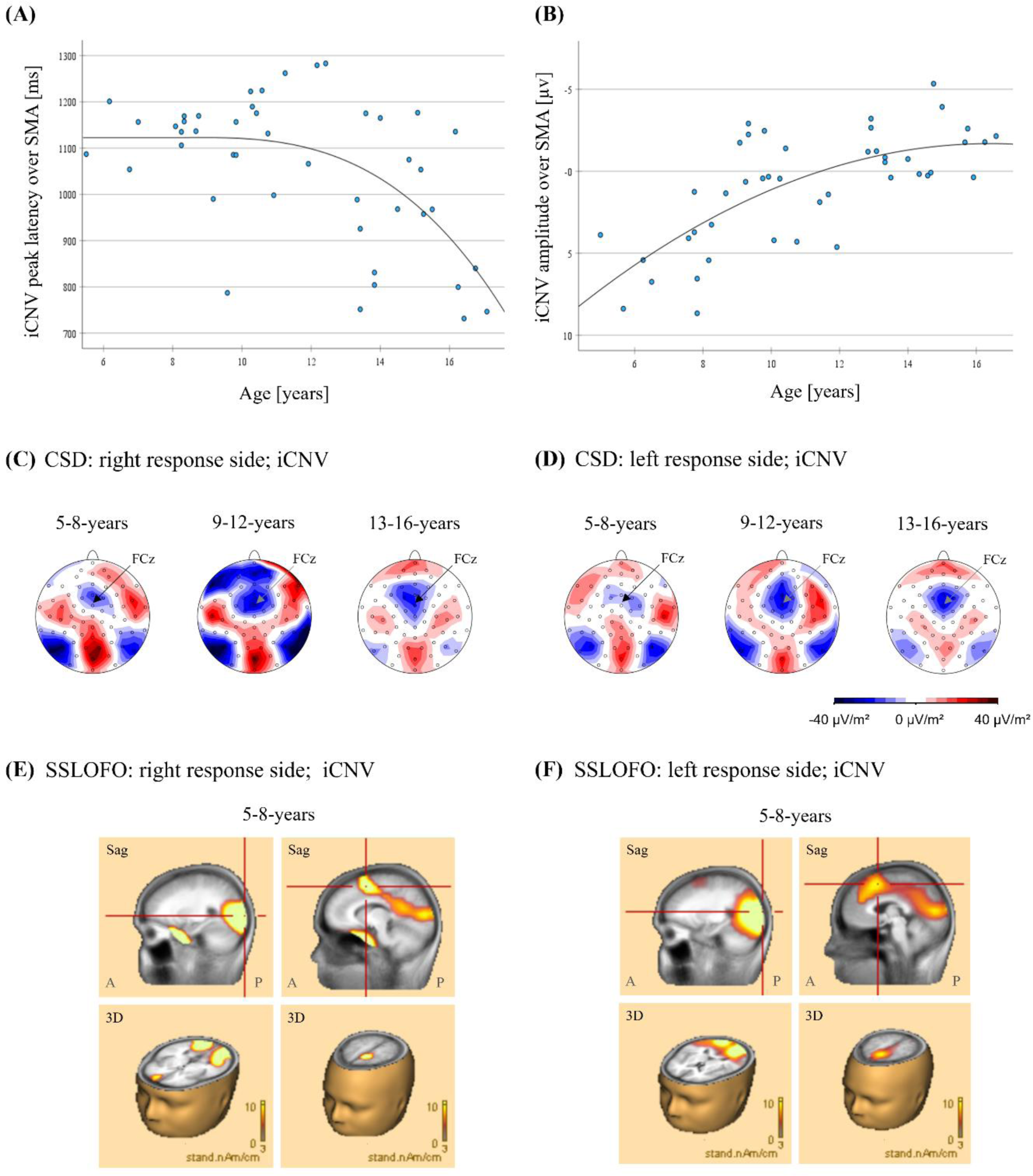
Developmental characteristics of the early CNV (iCNV) component. **(A)** Scatterplots of iCNV mean peak latencies and **(B)** amplitudes. Data points represent mean amplitudes for both response conditions and each subject over the mid-frontocentral area (Cz, FCz, FC1′, FC2′). The grey lines show the corresponding fitting of a curvilinear regression model (age^-1^), since age-related differences were more pronounced in younger subjects. **(C)** Average current source density (CSD) maps of the iCNV component for 5- to 8-year-old (left), 9- to 12-year-old (middle) and 13- to 16-year-old subjects (right), displayed for the right and **(D)** left response condition. CSD maps are scaled from -40 μV/cm^2^ (blue) to +40 μV/cm^2^ (red). **(E)** Image of current standardized shrinking LORETA-FOCUSS (SSLOFO) algorithm, calculated for right and **(F)** left response condition during iCNV. Images are displayed for 5- to 8-year-old subjects, to identify especially mid-frontocentral current sources in young subjects. Images showed a pronounced posterior (left images) as well as a mid-frontocentral cortical source (right images), related to the supplementary motor area. A: anterior side of brain; P: posterior side of the brain; Sag: Sagittal plane

#### 3.2.2 ERP amplitudes of iCNV component

The CNV topography of evoked cortical activity showed significant age-related developmental changes, most pronounced between the age range of 5- to 12-years. Figure 2 illustrates the stimulus-locked ERP waveforms for each group (for mean amplitudes with standard deviations see supplementary material, Table 1).

Comparing the evoked amplitudes of the early CNV component, the repeated measures ANOVA showed a highly significant interaction between scalp areas and age group (*F*(4, 76) = 6.76, *p* < .001).

Pairwise comparisons of estimated marginal means indicated significant differences in the activation of the mid-frontocentral area between the youngest age group of 5- to 8-year-olds (0.16 ± 2.78 µV) and the two older age groups of 9- to 12-year-olds (-3.66 ± 3.09 µV; *p* < .001) and 13- to 16-year-olds (-2.78 ± 2.03 µV; *p* = .01). Mid-frontocentral negativity of the youngest age group was shown not to differ significantly from baseline (see supplementary material, Table 1). In addition, no significant differences were observed when comparing the two older age groups. Centro-parietal scalp areas were found to exhibit no significant age effect during early movement planning.

To investigate correlation between the increasing mid-frontocentral activity and improved task performance, regression analyses were performed. The data showed a trend towards a curvilinear dependence between the mid-frontocentral activity and reaction time (RT^-1^: R^2^ = .115, F (1, 42) = 5.48, p = .024), however there was no significant correlation between mid-frontocentral activity and the performed errors (error rate: R^2^ = .104, F (1, 42) = 4.88, p = .033; significant at α < 0.0125).

#### 3.2.3 Alpha-event-related desynchronization during the iCNV component

Analysis of the alpha band event-related desynchronization (ERD) was conducted to investigate developmental differences of mu rhythm attenuation related to motor behavior planning and whether children aged 5- to 8-years showed early motor preparation, even when significant frontal ERP activity was not observed. Repeated measurement ANOVA revealed a main effect for the factor age group (*F*(2, 38) = 5.82, *p* = .006).

Pairwise comparison of estimated marginal means demonstrated an increase of alpha ERD with increasing age, as indicated by significant amplitude differences, observed between the youngest age group of 5- to 8-year-olds (-19.41 ± 4.4 µV) and the oldest age group of 13- to 16-years-olds (- 38.35 ± 3.89 µV; *p* =.008) as well as between the middle age group of 9- to 12-year-olds (- 24.72 ± 3.81 µV) and the oldest age group (*p* = .049). These findings suggest an increasing mu rhythm desynchronization associated with early movement planning throughout the examined age range.

In addition, there was a main effect of the scalp area (*F*(2, 76) = 45.87, *p* < .001) which was qualified by an interaction between movement side and scalp area (*F*(2, 76) = 25.84, *p* < .001). Alpha-ERD over contra- and ipsilateral centro-parietal areas but not over mid-frontocentral areas was shown to be highly significant for all age groups, as well as to be more pronounced over the contralateral hemisphere of the movement side (see Table 2). Pairwise comparison of estimated marginal means indicated that centro-parietal areas (right: - 32.17 ± 2.77 µV; left: - 32.32 ± 2.48 µV) showed significantly more desynchronization than the mid-frontocentral area (- 17.98 ± 2.33 µV, *p* < .001). Moreover, the movement side had a highly significant effect on alpha ERD over centro-parietal areas of both hemispheres (right, *p* < .001, lef: *p* < .001), showing a distinct lateralization to the contralateral movement side (see Figure 4C to 4F and supplementary material, Table 3). Thus, already 5- to 8-year-old children showed contralaterally lateralized alpha-ERD during early CNV (i.e., a selection of the required response side).

**Figure 4.**
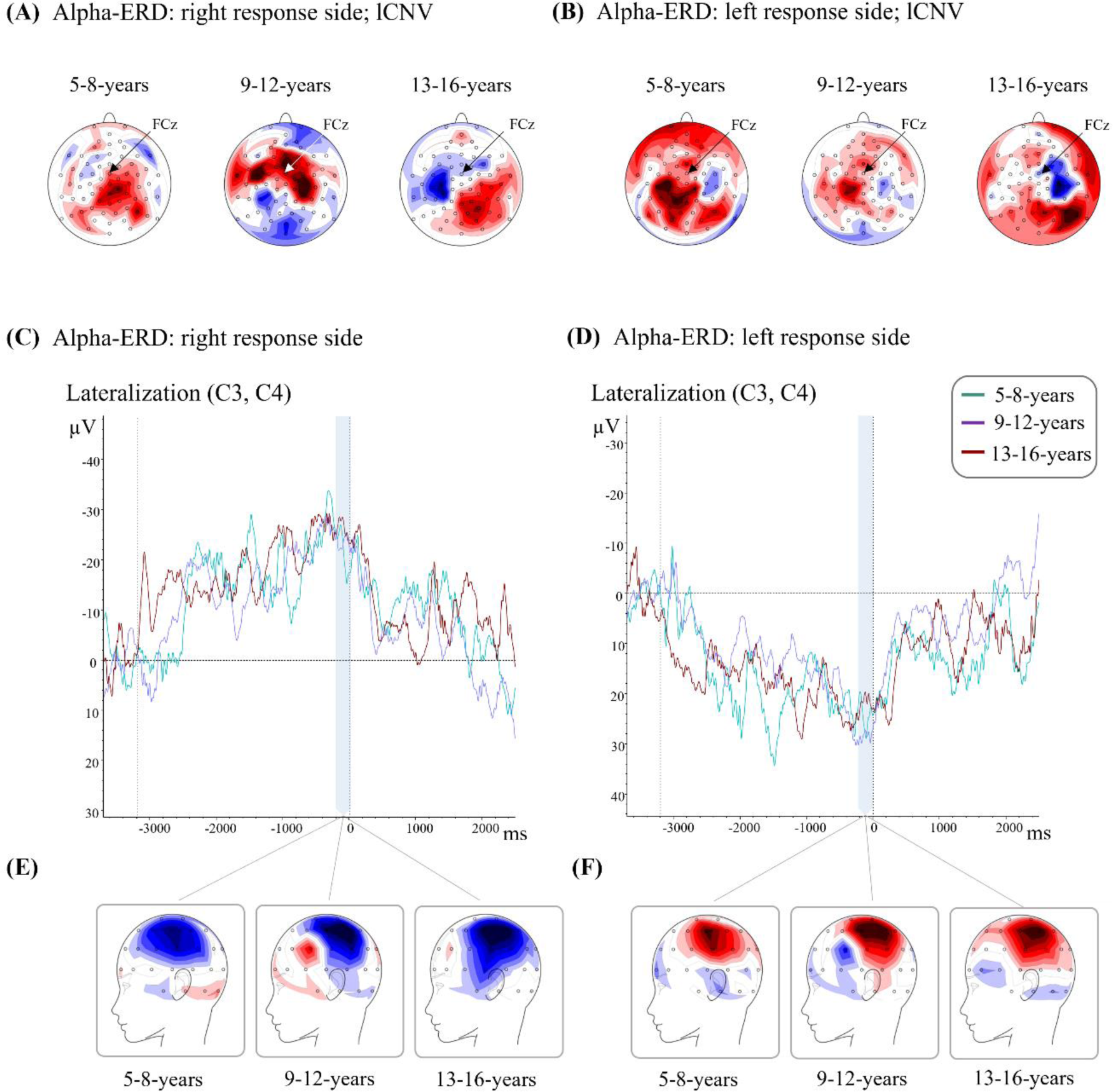
Developmental characteristics of the late CNV (lCNV) component. **(A)** Isopotential line maps of alpha event-related desynchronization (ERD) during lCNV, shown for right and **(B)** left response condition. Maps are displayed separately for average alpha ERD of 5- to 8-year-old (left), 9- to 12-year-old (middle) and 13- to 16-year-old subjects (right). Decrease of alpha power (ERD) is indicated in blue, increase of alpha power (event-related synchronization; ERS) is indicated in red. Scaling ranges from -30% to +30% in relation to baseline values. Ipsilateral positivity over central areas decreased with age, whereas contralateral negativity increased. **(C)** Lateralization (LRP) waveforms for the right and **(D)** left response condition, presented as average for 5- to 8-year-old subjects (green line), 9- to 13-year-old subjects (purple line) and 13- to 16-year-old subjects (red line). **(E)** LRP voltage maps of the contralateral hemisphere for the right response condition, **(F)** respectively for the ipsilateral hemisphere of the left response condition for 5- to 8-year-old (left), 9- to 12-year-old (middle) and 13- to 16-year-old subjects (right). Alpha-ERD showed an age-constant strong lateralization to the contralateral movement side during lCNV.

**Table 2.** Mean Alpha-ERD amplitude values [%] ± standard deviation for each CNV component. Significances (*p* ≤ 0.017) or trends towards significance (*p* < .33) are indicated for the respective values.

| *Alpha-ERD of CNV Component* | *Age Group* | *SMA [%]* | *M1 right [%]* | *M1 left [%]* |
| --- | --- | --- | --- | --- |
| *iCNV right response condition* | 5- to 8-year-old | $-6.27 \pm 16.54$ | $-16.24 \pm 23.57$<br>$t(-2.38); p=.03$ | $-29.32 \pm 22.05$<br>$t(-4.61); p < .001$ |
| | 9- to 12-year-old | $-12.51 \pm 12.79$<br>$t(-4.03); p < .001$ | $-23.99 \pm 18.08$<br>$t(-5.47); p < .001$ | $-34.27 \pm 16.6$<br>$t(-8.51); p < .001$ |
| | 13- to 16-year-old | -33.06 ± 21.39<br>$t(-5.78); p < .001$ | -35.79 ± 25.53<br>$t(-5.25); p < .001$ | -51.99 ± 19.55<br>$t(-9.95); p < .001$ |
| *iCNV left response condition* | 5- to 8-year-old | -13.48 ± 18.88<br>$t(-2.47); p = .03$ | -34.59 ± 14.83<br>$t(-8.08); p < .001$ | -22.21 ± 16.84<br>$t(-4.57); p < .001$ |
| | 9- to 12-year-old | -17.14 ± 12.39<br>$t(-5.71); p < .001$ | -37.33 ± 15.8<br>$t(-9.74); p < .001$ | -28.02 ± 15.27<br>$t(-7.57); p < .001$ |
| | 13- to 16-year-old | -29.34 ± 19.5<br>$t(-5.63); p < .001$ | -46.64 ± 22.08<br>$t(-7.9); p < .001$ | -32.74 ± 21.71<br>$t(-5.64); p < .001$ |
| *lCNV right response condition* | 5- to 8-year-old | 9.22 ± 22.49 | 12.94 ± 22.65<br>$t(1.98); p = .05$ | -0.88 ± 20.88 |
| | 9- to 12-year-old | 12.94 ± 20.23<br>$t(2.64); p = .02$ | 24.51 ± 36.32<br>$t(2.78); p = .01$ | -9.23 ± 20.49<br>$t(-1.86); p = .08$ |
| | 13- to 16-year-old | -4.44 ± 22.27 | 11.21 ± 33.57 | -17.94 ± 23.53<br>$t(-2.85); p = .01$ |
| *lCNV left response condition* | 5- to 8-year-old | 6.91 ± 20.56 | -5.44 ± 15.65 | 23.05 ± 22.12<br>$t(3.61); p = .004$ |
| | 9- to 12-year-old | 5.39 ± 17.78 | -6.27 ± 16.57 | 14.44 ± 29.22<br>$t(2.04); p = .06$ |
| | 13- to 16-year-old | -0.78 ± 17.74 | -16.39 ± 21.64<br>$t(-2.84); p = .01$ | 1.87 ± 28.42 |
| *PINV right response condition* | 5- to 8-year-old | -14.18 ± 17.43<br>$t(-2.82); p = .01$ | -28.78 ± 19.72<br>$t(-5.06); p < .001$ | -33.39 ± 21.89<br>$t(-5.28); p < .001$ |
| | 9- to 12-year-old | -24.46 ± 16.12<br>$t(-6.25); p < .001$ | -35.88 ± 19.22<br>$t(-7.7); p < .001$ | -42.68 ± 20.93<br>$t(-8.41); p < .001$ |
| | 13- to 16-year-old | -38.86 ± 25.46<br>$t(-5.71); p < .001$ | -46.71 ± 25.26<br>$t(-6.92); p < .001$ | -53.19 ± 23.43<br>$t(-8.35); p < .001$ |
| *PINV left response condition* | 5- to 8-year-old | -18.07 ± 20.89<br>$t(-2.98); p = .01$ | -37.56 ± 19.86<br>$t(-6.55); p < .001$ | -23.38 ± 25.36<br>$t(-3.19); p = .009$ |
| | 9- to 12-year-old | -24.37 ± 18.82<br>$t(-5.34); p < .001$ | -42.45 ± 20.31<br>$t(-8.61); p < .001$ | -43.86 ± 19.72<br>$t(-9.17); p < .001$ |
| | 13- to 16-year-old | -39.88 ± 22.26<br>$t(-6.7); p < .001$ | -54.66 ± 24.5<br>$t(-8.35); p < .001$ | -48.49 ± 27.31<br>$t(-6.64); p < .001$ |

#### 3.2.4 Current source density (CSD) of iCNV component

The qualitatively distinct pattern observed during iCNV in young subjects could also be evoked by strong, overlapping P300 activity masking the anticipated activity that was shown in older subjects. To investigate activity over mid-frontocentral areas in younger subjects more precisely and to reduce the influence of overlapping P300 activity, we conducted Current Source Density (CSD) analysis.

To assess age effects, we conducted a repeated measures ANOVA, which indicated an interaction between scalp area and age group (*F*(4, 76) = 3.13, *p* = .019), suggesting age-related topographical maturation. Pairwise comparison of estimated marginal means revealed age-related activity differences over the centro-parietal scalp areas between the youngest (right: -29.48 ± 9.12 µV; left: -23.72 ± 9.96 µV) and the oldest age group (right: 4.09 ± 8.06 µV*, p* = .026; left: 10.0± 8.8 µV*, p* = .045), showing decreasing negativity over left and right centroparietal areas with age. However, CSD map topographies indicated that the topographic maximum related to these current sinks was over occipito-temporal areas, so we did not interpret this activation any further with respect to motor processes.

More importantly, the mid-frontocentral area exhibited no significant differences between the age groups. Further examination of CSD map topographies (Figure 3B, 3C) revealed that even children in the age range of 5- to 8-years exhibited a small current sink over mid-frontocentral areas (-10.92 ± 32.9 µV) which was however not significant (*t*(11) = - 1.15, *p* = .27). Figure 3B and 3C illustrate that in younger subjects the positivity of the P3-complex and an occipito-temporal negativity over visual cortical areas dominated the CSD maps. Conversely, in older subjects, mid-frontocentral negativity became more prominent.

#### 3.2.5 Source analysis (SSLOFO) of iCNV component

SSLOFO analysis, an inverse algorithm to precisely reconstruct underlying sources, was applied to identify closely spaced cortical generators of the mid-frontocentral activity during iCNV of young subjects. Consistent with the findings from CSD analysis (overlapping P3-complex, occipito-temporal current sinks), SSLOFO analysis confirmed the presence of a prominent posterior source in young subjects during early movement planning. However, source analysis indicated that already in young subjects from the age of 5 years, the supplementary motor area is likely to contribute, at least partly, to early negativity observed over mid-frontocentral areas (Figure 3D and 3E).

### 3.3 Late motor preparation (late contingent negative variation)

#### 3.3.1 ERP amplitudes of lCNV component

To investigate maturational processes related to direct motor preparation, we focused on the late component of the CNV paradigm. The lCNV topography showed pronounced differences between the age groups (see Figure 2C and 2D and supplementary material, Table 1). Children aged 5 to 8 years displayed more widespread low ipsilateral negativity, most pronounced over centroparietal areas. In contrast, older subjects (9- to 16-year-olds) exhibited a more centrally localized negativity over mid-frontocentral and central areas, with decreasing activity over ipsilateral centro-parietal areas with age.

The repeated measurement ANOVA revealed significant interactions between scalp area and movement side (*F*(2, 76) = 3.59, *p* = .034) and between scalp area and age group (*F*(4, 76) = 3.63, *p* = .009). Pairwise comparison of estimated marginal means showed that the movement side had a significant effect on the left centroparietal scalp area (right hand movements: - 0.36 ± 0.32 µV; left hand movements: - 1.48 ± 0.3 µV, *p* = .017), with more pronounced ipsilateral negativity observed for the left movement side. The right centroparietal scalp area was observed not to be significantly influenced by the movement side.

Furthermore, significant amplitude differences over the mid-frontocentral area were observed between the youngest age group of 5- to 8-year-olds (-0.74 ± 0.73 µV) and the middle age group of 9- to 12-year-olds (-3.51 ± 0.58 µV; *p* = .02). There was also a trend towards more midfronto-central negativity for the oldest age group of 13- to 16-year-olds (- 3.04 ± 0.64 µV; *p* = .067) compared to 5- to 8-year-olds. Centro-parietal areas of both the contralateral and ipsilateral hemisphere did not show significant maturational changes.

#### 3.3.2 Alpha-Event-related desynchronization of lCNV component

To investigate oscillatory modulation related to movement preparation and cortical pre-activation prior to a movement, we analyzed alpha ERD during lCNV.

Repeated measurement ANOVA showed a significant interaction between movement side and scalp area (*F*(2, 76) = 25.59, *p* < .001). Pairwise comparison of the estimated marginal means revealed that alpha ERD over centroparietal areas were highly influenced by the movement side (*p* <.001).

Despite pronounced maturational differences of alpha power changes displayed in the related maps and data (Figure 4A and 4B, Table 2), ANOVA analysis showed a trend towards differences between the age groups (*F*(2, 38) = 2.84, *p* = .071). To investigate the data in more detail, we performed a regression analysis over centroparietal motor areas. Regression analysis confirmed a strong trend towards a developmental trajectory with decreasing alpha band synchronization over ipsilateral motor areas and increasing desynchronization over contralateral motor areas (Age; ipsilaterally decreasing synchronization: R^2^ = .109, F (1, 42) = 5.16, *p* = .028; contralaterally increasing desynchronization: R^2^ = .08, F (1, 42) = 3.77, *p* = .059; significant for α < .025) (for details see supplementary material, Figure 1).

Interestingly, an analysis of the lateralization of alpha-ERD further illustrated that a constant degree of lateralization was maintained during development (Figure 4C to 4D).Topographic distribution of the lateralized alpha-ERD during lCNV (Figure 4E and 4F) as well as lateralized alpha-ERD amplitude analysis (*F*(2, 38) = 0.014, *p* = .99) confirmed no significant lateralization differences between the analyzed age groups, despite apparent differences in the contra- and ipsilateral induced alpha power (see Figure 4A, 4B for the time course and topography of lateralized activation).

### 3.4 Response postprocessing and evaluation (post imperative negative variation, PINV)

#### 3.4.1 Latency of PINV component

To investigate maturational changes of response evaluation processes, we analyzed cortical activity related to motor post-processing (PINV) within a CNV paradigm. Repeated measurement ANOVA showed a significant interaction between movement side, the scalp area and the age group (*F*(3.86, 73.3) = 2.75, *p* = .036).

Pairwise comparison of estimated marginal means indicated that, for the right movement side, negativity over the right ipsilateral hemisphere arose with an increased latency in the oldest age group (1171.18 ± 59.59 ms), compared to the youngest (897.47 ± 67.42 ms, *p* = .013) and the middle age group (865.71 ± 58.43 ms, *p* = .002). For the left-hand movement, no significant latency differences were observed.

#### 3.4.2 ERP amplitudes of PINV

PINV topography varied strongly between the age groups (see Figure 2). Younger subjects displayed maximum negativity over centro-parietal areas, while in older subjects pronounced negativity was shifted towards more mid-frontocentral areas. The repeated measurement ANOVA indicated a statistically significant interaction between scalp area and age group (*F*(4, 76) = 4.12, *p* = .004) and between the movement side and scalp area (*F*(2, 76) = 8.9, *p* < .001).

Pairwise comparison of estimated marginal means showed that the movement side had a significant effect on the right (*p* = .009) and left centro-parietal scalp area (*p* = .001), with more pronounced negativity observed over the contralateral hemisphere.

Maturational influences were found over the left centro-parietal scalp area, with a trend towards amplitude differences between the youngest (- 3.56 ± 0.77 µV) and the oldest age group (- 1.03 ± 0.68 µV; *p* = .054), showing decreasing PINV amplitude over the left-centroparietal area with age. Furthermore, negativity shifted from centro-parietal scalp areas in young subjects to mid-frontocentral areas in older subjects (see supplementary material, Table 1).

#### 3.4.3 Alpha-Event-related desynchronization of PINV component

To study maturational changes of alpha band oscillation related to response evaluation, we analyzed alpha-ERD during PINV. Desynchronization of alpha power was present for all age groups over mid-frontocentral as well as over centro-parietal areas, and most pronounced in older subjects (Table 2).

ANOVA analysis showed an interaction between movement side and scalp area (*F*(2, 76) = 9.23, *p* < .001). Pairwise comparison of the estimated marginal means indicated that alpha desynchronization was more pronounced over contralateral centro-perietal scalp areas, especially for the right hemisphere (*p* < .001).

Furthermore, there was a main effect of age group (*F*(2, 38) = 5, *p* = .012) as well as a trend towards an interaction between the scalp area and the age group (*F*(4, 76) = 2.23, *p* = .074). In total, alpha desynchronization was less pronounced in younger subjects aged 5- to 8-years (-24.71 ± 5.33 %) than in older subjects aged 13- to -16 years (- 46.61± 4.72%; *p* = .012). Regarding maturational differences of the separated scalp areas, negativity over mid-frontocentral areas was more pronounced in the oldest age group (- 39.0± 4.6%), compared to the youngest (- 15.05± 5.21%; p = .004) and the middle age group (- 21.51± 4.51%; p = .03).

## 4 Discussion

Brain maturational processes and underlying neuronal mechanisms involved in the transition from an immature child brain to a purposive adult brain are complex and difficult to identify. Our data provide evidence of pronounced cerebral maturation related to attention allocation, motor preparation and movement evaluation during childhood and adolescence. The study compared behavioural task performance and cortical activation evoked and induced by a contingent negative variation (CNV) paradigm with a directional warning cue. We collected EEG data of subjects aged of 5- to 16-years using a 64 equidistant electrode array.

Our findings provide evidence that enhanced SMA activity plays a critical role in increasing efficiency of motor behaviour during cortical maturation. Besides poorer task performance, young subjects displayed qualitatively different cortical activation patterns during the stages of movement planning, preparation and post-processing compared to older subjects. SMA activity showed a developmental increase during early movement selection and subsequent motor preparation maintenance as well as during motor post-processing.

ERP data of young subjects aged 5- to 8-years revealed no significant, however small beginning activation of SMA related cortical areas during processes of early movement selection (iCNV). A pronounced contralateral alpha power reduction of motor related scalp areas, accordingly an increase of cortical activity, was observed for all age stages. This finding is consistent with an early alerting of sensory and motor regions corresponding to the required movement side (Babiloni et al., 1999).

During late movement preparation (lCNV), alpha power showed a trend towards a maturational transition from ipsilateral synchronization (i.e. inhibition; deactivation) to contralateral desynchronization (i.e. disinhibition; pre-activation) within motor areas. These results support the hypothesis of a developmental shift from a reactive to a proactive motor control (Chevalier et al., 2014). In contrast to other studies (Bender et al., 2005; Flores et al., 2009) adolescent subjects showed no evoked contralateral motor area negativity during late movement preparation. The observed differences might be induced by varying task characteristics, e.g. early selection of the movement side. Pronounced pre-activation of effector specific motor areas (Dirnberger et al., 2003) seems to develop later in more complex processes of motor preparation.

Movement post-processing (PINV) was characterized by a trend towards age-related reduction of evoked negativity over contralateral motor areas and a simultaneous increase of SMA related negativity. Since pronounced activation of motor areas during PINV(Thiemann, 2010)) is associated with uncertainty of the performed movement (Bender et al., 2006), it suggests that reduced motor control in young children is reflected in enhanced motor post-processing.

### 4.1 Behavioural performance

Reaction times as well as performed errors improved with age, as typically observed in other studies (Bender et al., 2005; Bucsuházy & Semela, 2017; Kiselev et al., 2009). Motor improvement was characterized by shorter reaction times and reduced false alarms or wrong responses in older subjects. The observed developmental differences had an influence on the processing speed as well as on the ability to restrain motor behaviour. Improved action control developed curvilinearly in the age range of 5 to 16 years, with most pronounced maturation in the youngest age group of 5- to 8-year-old children.

### 4.2 Maturation of orienting response (early contingent negative variation)

The early or initial CNV component (iCNV) is associated with orienting processes as a response to a warning stimulus in a CNV paradigm (Weerts & Lang, 1973). According to Birbaumer et al. (1990), the iCNV reflects prefrontal activation that regulates activation of more posterior motor areas necessary for movement execution.

The early component was investigated to study the impact of cerebral maturation on the orienting response to an informative external stimulus, including attentional processes and motor program selection. ERP analysis revealed immature iCNV topographies until late childhood and early adolescence. Electrophysiological data showed a pronounced maturation characterized by increasing early mid-frontocentral negativity through the investigated age range. In agreement with other studies using data of a reduced number of electrodes, young children showed only minor frontal negativity during orienting response (Jonkman et al., 2003; Segalowitz & Davies, 2004), i.e., activation of cortical areas related to the supplementary motor area (SMA) and anterior cingulate cortex (ACC) (Flores et al., 2009; Jonkman et al., 2003). The early frontal negativity stabilized during post-pubertal adolescence, reaching strength of activation comparable to adult subjects (Tian et al., 2019; Weisz et al., 2002). Furthermore, we observed a consistent shift in iCNV latencies, with mid-frontocentral areas being recruited earlier in older subjects, suggesting a faster processing of the warning stimulus.

Besides small frontal negativity, young children showed enhanced processing of the directional warning cue over posterior-temporal and occipital areas related to visual cortices (Bender et al., 2010; Hecht et al., 2016) that caused a partial masking of small SMA activity in the ERP analysis. Van Leeuwen et al. (1998) observed similar early topographies and found posterior sources for this so called CNV/P3 complex. It is likely that other studies reporting contradictory decreasing mid-frontocentral negativity during iCNV with age (Bender et al., 2004) as well as reduced iCNV latencies in younger children (Flores et al., 2009) observed rather fronto-central negativity that was dependent on extended stimulus processing related to a posterior source than true frontal iCNV negativity associated with SMA activity. Enhanced activity reflected in a more pronounced CNV/P3 complex could be explained by an increased effort to process relevant task information. The orienting reaction is thus dominated by sensory post-processing and resource allocation in young children, while more frontal activation with decreasing latencies becomes a dominant part in adolescent subjects.

Since young children reacted with more impulsive motor responses and showed an increased error rate, posterior networks recruited during movement preparation seem to be less efficient in attention regulation and action control. Precipitated movements might be caused by inefficient motor inhibition or by reduced efficiency of attention allocation due to an increased effort in target selection processes. The observed maturational shift is consistent with described strategy changes from a posterior, stimulus driven orienting network in young children to an anterior, top-down attention network in adolescent subjects (Padilla et al., 2014; Smith et al., 2011).

Cortical source analysis of iCNV data using sSLOFO confirmed that the mid-frontocentral negativity is likely to originate from the supplementary motor area (SMA). It was suggested that the SMA recruits and sustains networks that are activated during subsequent preparatory processes related to the lCNV (Gomez et al., 2003). In other studies, it was shown that the iCNV arises from the SMA and ACC and that the cortical areas were simultaneously coactivated (Gomez et al., 2003; Lee et al., 1999). Lee et al. (1999) observed that subregions of the SMA were recruited with different temporal profiles and that early stages of motor processing were associated with the activation of anterior SMA parts related to the pre-SMA. The pre-SMA is characterized by extensive pre-frontal connectivity (Luppino et al., 1993) and thought to be involved, inter alia, in maintaining working memory (Pollmann & Yves von Cramon, 2000) as well as motor program selection and movement intention (Lau et al., 2004). Nachev et al. (2007) performed a movement study in a patient with a rare lesion involving the pre-SMA and reported increased reaction times as well as inhibitory deficits. Other studies confirmed that the SMA mediates motor inhibition (Chen et al., 2010; Toma et al., 1999). These results suggest that the pre-SMA exhibits a critical role in action control and indicate that pre-SMA immaturity might be a primary cause for less efficient movement performances in young children. However, it is not possible to exclude an indirect correlation between task performance and SMA activation due to parallel age effects. Therefore, a direct functional relevance of the increased SMA activation in improving movement performance cannot be investigated based on the observed data.

Brain oscillatory activity was analyzed to gain insight into maturational changes of movement-related power modulation. Mu-rhythm desynchronization is observed for cortical areas involved in processes of movement preparation, planning and execution (Leocani et al., 1997; Pfurtscheller & Lopes da Silva, 1999). During cognitive processing or attention allocation processes, alpha power is reduced and facilitates resources to be allocated to task-relevant processing. Cortical regions that are related to task-irrelevant and potentially disruptive processes exhibit mu-rhythm synchronization, i.e. inhibition of underlying areas (Klimesch et al., 2007). The results showed lateralized mu-rhythm desynchronization over contralateral central areas during iCNV for all age groups. This suggests that contralateral motor areas were activated during early processes of movement side selection shortly after the presentation of the informative warning cue. Given that the early motor preparation in young children was already characterized by a lateralized activation of contralateral motor regions, it suggests that also young children exhibit motor related preparatory processes. Moreover, the results confirm that motor program selection corresponds to early stages of motor preparation, as reflected by contralateral motor area activation.

Compared to evoked cortical potentials, Mu-rhythm desynchronization reflects more general sensorimotor network pre-activation (C. Babiloni et al., 1999; Defebvre et al., 1994). In contrast to evoked potentials, differences of early mu rhythm desynchronization over mid-frontocentral areas were most pronounced between the middle (9- to 12-year-olds) and the oldest age group (13- to 16-year-olds), which suggests a delayed development of preparatory alpha band power in relation to evoked cortical activity.

### 4.3 Maturation of motor cortical pre-activation (late contingent negative variation)

The late component of a CNV has been suggested to represent sensory and motor pre-activation of resources needed for effective task-specific performance (Gaillard, 1978). Midfronto-central negativity increased with age during response preparation (lCNV). Previous results confirm the continuous development of preparatory movement networks and maturation of the motor system into young adulthood (Bender et al., 2002; Jonkman, 2006; Killikelly & Szucs, 2013; Thillay et al., 2015). Significant lCNV activity indicates that adolescent subjects used proactive control strategies to perform more efficient task-related movements.

The late CNV component is known to be prominent over contralateral hemispheres (Dirnberger et al., 2003; van der Lubbe et al., 2000; Wauschkuhn et al., 1997). However, in contrast to developmental studies using simple CNV paradigms (Bender et al., 2005), the ERP data showed transient contralateral negativity only during early motor preparation, which might be related to early motor program selection. Late contralateral activity over pre- and primary motor areas, which is usually observed in adolescent and young adult subjects (Bender et al., 2005; Flores et al., 2009), was not confirmed in our adolescent sample. Since the activation patterns of the analyzed adolescent subjects (up to the age of 16 years) were therefore qualitatively different from activation patterns typically observed in adult subjects (Gomez et al., 2003), results suggest an ongoing development of cortical areas related to preparatory activity.

Task characteristics and paradigm complexity have an influencing effect on CNV waveforms, topographies, and amplitudes. The warning cue of the CNV paradigm indicated the required movement side and was therefore more complex than simple paradigms used in other studies without varying warning stimulus information or movement side. Since an additional component was added to our paradigm, task-related preparatory processes required higher levels of executive motor control. Development of sustained lateralized negativity over the contralateral central areas may be delayed with respect to studies of simple CNV paradigms. We suggest that effective motor preparation of more complex movement tasks is achieved at a late stage of maturation and that movement-related cortical development is not completed until late adolescence or young adulthood.

As in other studies, it was shown that the SMA not only played a major role in processes related to the early CNV component but also to late CNV processes (Nagai et al., 2004). Regarding the adolescent subjects in our sample, the topographic analysis revealed a transition from frontal to more central negativity when comparing early and late CNV stages. This change in topography could be associated with a shift from increased pre-SMA activation during early pre-processing (iCNV) to higher activity of more caudal SMA parts (SMAc) and premotor areas during late motor pre-activation (lCNV). Whereas the anterior part of the SMA is associated with early pre-processing stages related to motor program planning, the more caudal part is suggested to be involved in later stages related to movement initiation and execution (Lee et al., 1999). Connectivity studies of the SMA confirmed this assumption. In contrast to pre-SMA, which is connected to prefrontal areas (Wang et al., 2005), SMAc has direct connections to the primary motor cortex (Luppino et al., 1993).

Time-frequency analysis revealed that alpha-ERD (excitation) of the contralateral hemisphere linearly increased with age during late movement preparation and planning (lCNV), whereas alpha-ERS (inhibition) of the ipsilateral side decreased. However, lateralization analysis of the alpha power band showed no differences in lateralization strength between young children and adolescent subjects, as already observed during iCNV. Since synchronized activity is related to deactivation of corresponding brain areas (Lopes da Silva, 2006; Pfurtscheller, 2001), the results suggest that processes related to alpha band oscillation of motor networks were shifted from inhibition of ipsilateral motor-related brain areas to pre-activation of contralateral motor-related areas. These findings sustain the hypothesis of a maturational shift from a reactive to a proactive motor control (Chevalier et al., 2014). Furthermore, age-consistent alpha band lateralization showed that small SMA activity observed for young subjects was not evoked due to low motor preparation or a lack of motivation but due to qualitatively different preparatory processes.

### 4.4 Maturation of movement processing (post imperative negative variation)

The PINV is associated with movement evaluation processes and is suggested to represent uncertainty in task performance (Bender et al., 2006; Klein et al., 1996). Reliance of the PINV on response execution was confirmed due to its correlation with response execution timing rather than the timing of external stimuli (Bender et al., 2004). EEG studies showed enhanced PINV amplitudes in children and adult subjects when they had to perform non-controllable tasks (Kathmann et al., 1990; Yordanova et al., 1997).

ERP data showed a developmental shift from pronounced negativity over contralateral central areas to negativity over more frontocentral areas during performance evaluation. The PINV did not depend on preparatory CNV components since topographical distribution differed significantly, as shown in previous studies (Bender et al., 2004; Bender et al., 2005). Since performance of younger subjects was associated with a lack of control, high PINV amplitudes might reflect controllability of task-related movements. Moreover, the results suggest that young children were more likely to be uncertain about their response performance, reflected in an increased compensatory effort and enhanced evaluation processes. Future studies should address the question whether increased motor post-processing supports motor learning in children.

As observed for evoked potentials, alpha band desynchronization increased with age over mid-frontocentral areas during response evaluation. These results indicate that mid-frontocentral areas associated with SMA activity are increasingly recruited with age during movement post-processing and motor maturation. Moreover, evoked PINV topographies showed most pronounced maturational changes for the youngest and the middle age group, whereas alpha-ERD was characterized by pronounced developmental differences between the middle and the oldest age group, indicating an ongoing maturation of alpha band oscillation until late adolescence or early adulthood.

### 4.5 Limitations

Based on the presented results, it is not possible to identify physiological mechanisms that underly the observed changes in cortical activity and alpha band oscillation. It is likely that the observed maturation depends, inter alia, on developmental changes in functional connectivity that induce altered network dynamics (Brookes et al., 2018). Future studies should include multi-modal comparisons (fMRI) and connectivity analyses to investigate brain development more precisely. Moreover, studies of developmental changes should include an analysis of longitudinal examinations and not be based exclusively on cross-sectional data. The investigation of cortical maturation on an individual, intra-subject level allows the generation of more reliable and accurate information.

## 5 Conclusion

Our data showed pronounced maturational differences in cortical movement preparation and post-processing between child- and adolescent subjects. Behavioural results indicated less-efficient action pre-processing and a lack of inhibitory control, probably due to incomplete frontal lobe maturity in children. Activation of mid-frontocentral areas related to the supplementary motor area during movement preparation and evaluation became more prevalent with age. Alpha band power indicated developmental progress from inhibition of ipsilateral motor areas in young children to increased pre-activation of contralateral motor areas in adolescents. Based on our data, evoked cortical activation is likely to develop earlier than alpha band oscillatory activity. The reported results supported the hypothesis of a developmental shift from a reactive to a proactive control and an immaturity of supplementary-, pre- and primary motor areas until late adolescence or early adulthood.

## Abbreviations

ACC: Anterior cingulate cortex
CNV: Contingent negative variation
CSD: Current source density
EEG: Electroencephalogram
EHI: Edinburgh Handedness Inventory
ERD: Event-related desynchronization
ERP: Event-related potential
ERS: Event-related synchronization
ICA: Independent component analysis
iCNV: initial or early CNV
lCNV: late CNV
LORETA: Low Resolution Brain Electromagnetic Tomography
LRP: Lateralized readiness potential
M1: primary motor cortex
MRP: Motor evoked potential
PINV: Post-imperative negative variation
SMA: Supplementary motor area
SSLOFO: Current Standardized shrinking LORETA-FOCUSS

## 5.2 Acknowledgements

We thank all our participants as well as Elena Borovik, Sally Kufall and Amelie Busch for their assistance in data acquisition.

## 5.3 Authors contribution

J.S.: Acquisition of data; data analysis; visualization of data; interpretation of results; writing of the manuscript. T.H.: Acquisition of data; proofreading of the manuscript. K.K.: Study conception and design; editing and critical writing revision. S.B.: Study conception and design; supervision of analysis and interpretation of data; editing and critical writing-revision. All authors discussed the results, contributed to the final manuscript and gave final approval of the version to be published.

## 5.4 Ethics approval statement

This study was conducted in accordance with the local legislation and institutional requirements. The research protocol was reviewed and approved by the Ethics Committee of the University of Cologne and the Ethics Committee of the University of Aachen.

