## Supplementary Material for "Mapping Motor Preparation in the Developing Brain: Insights from Contingent Negative Variation and Event-related Mu Rhythm Modulation"

### 1 Supplementary material

Table 1. Mean ERP amplitude values [µV] ± standard deviation for each CNV component. Significances (*p* ≤ 0.17) or trends towards significance (*p* < .33) are indicated for the respective values.

| Evoked activity of CNV Component | Age Group | SMA [µV] | M1 right [µV] | M1 left [µV] |
| --- | --- | --- | --- | --- |
| iCNV right response condition | 5- to 8-year-old | 0.28 ± 2.33 | -0.44 ± 2.9 | -0.11 ± 2.37 |
|  | 9- to 12-year-old | -3.87 ± 2.89  *t*(-5.52); *p* < .001 | -0.66 ± 2.08 | -1.73 ± 2.44  *t*(-2.93); *p =* .01 |
|  | 13- to 16-year-old | -2.98 ± 2.32  *t*(-4.99); *p* < .001 | 0.4 ± 1.11 | -0.42 ± 1.86 |
| iCNV left response condition | 5- to 8-year-old | 0.04 ± 3.24 | -0.8 ± 3.05 | -1.15 ± 1.38  *t*(-2.88); *p =* .02 |
|  | 9- to 12-year-old | -3.45 ± 3.29  *t*(-4.33); *p* < .001 | 0.02 ± 2.34 | -1.11 ± 2.77 |
|  | 13- to 16-year-old | -2.58 ± 1.75  *t*(-5.73); *p* < .001 | -0.21 ± 1.5 | 0.06 ± 2.11 |
| lCNV right response condition | 5- to 8-year-old | -0.4 ± 3.21 | -0.6 ± 2.83 | -0.59 ± 1.59 |
|  | 9- to 12-year-old | -3.66 ± 2.66  *t*(-5.67); *p* < .001 | -2.49 ± 2.07  *t*(-4.96); *p* < .001 | -0.17 ± 2.08 |
|  | 13- to 16-year-old | -3.3 ± 2.13  *t*(-6); *p* < .001 | -1.23 ± 1.37  *t*(-3.48); *p =* .004 | -0.31 ± 2.16 |
| lCNV left response condition | 5- to 8-year-old | -1.32 ± 3.61 | -0.4 ± 4.35 | -1.7 ± 1.87  *t*(-3.15); *p =* .009 |
|  | 9- to 12-year-old | -3.52 ± 3.62  *t*(-4.02); *p* < .001 | -1.05 ± 1.52  *t*(-2.84); *p* = .01 | -1.31 ± 1.91  *t*(-2.84); *p* = .01 |
|  | 13- to 16-year-old | -2.74 ± 1.53  *t*(-6.96); *p* < .001 | -1.74 ± 1.49  *t*(-4.54); *p* < .001 | -1.66 ± 1.87  *t*(-3.43); *p* = .004 |
| PINV right response condition | 5- to 8-year-old | -0.25 ± 2.94 | -1.32 ± 4.29 | -4.13 ± 3.01  *t*(-4.75); *p* < .001 |
|  | 9- to 12-year-old | -2.73 ± 3.04  *t*(-3.7); *p* = .002 | -1.57 ± 3.16  *t*(-2.05); *p* = .057 | -2.53 ± 3.23  *t*(-3.23); *p* = .005 |
|  | 13- to 16-year-old | -3 ± 2.18  *t*(-5.35); *p* < .001 | 0.15 ± 1.97 | -1.49 ± 1.1  *t*(-5.26); *p* < .001 |
| PINV left response condition | 5- to 8-year-old | -1.48 ± 3.14 | -2.79 ± 4.32  *t*(-2.23); *p =* .05 | -3.02 ± 2.95  *t*(-3.54); *p =* .005 |
|  | 9- to 12-year-old | -3.15 ± 3.66  *t*(-3.55); *p* = .003 | -2.7 ± 2.08  *t*(-5.35); *p* < .001 | -1.21 ± 3.66 |
|  | 13- to 16-year-old | -2.49 ± 1.65  *t*(-5.86); *p* < .001 | -1.24 ± 1.86  *t*(-2.58); *p* = .02 | -0.56 ± 1.81 |

Table 2 Mean ERP latency values [ms] ± standard deviation for each CNV component.

| Latencies of evoked activity of CNV Component | Age Group | SMA [ms] | M1 right [ms] | M1 left [ms] |
| --- | --- | --- | --- | --- |
| iCNV right response condition | 5- to 8-year-old | 1116 ± 115 | 951 ± 155 | 1010 ± 230 |
|  | 9- to 12-year-old | 1057 ± 177 | 965 ± 223 | 1160 ± 192 |
|  | 13- to 16-year-old | 982 ± 161 | 1075 ± 183 | 1112 ± 163 |
| iCNV left response condition | 5- to 8-year-old | 1136 ± 73 | 1091 ± 263 | 1108 ± 137 |
|  | 9- to 12-year-old | 1132 ± 182 | 1155 ± 215 | 1064 ± 149 |
|  | 13- to 16-year-old | 941 ± 294 | 1072 ± 205 | 998 ± 171 |
| PINV right response condition | 5- to 8-year-old | 1102 ± 203 | 918 ± 276 | 1023 ± 112 |
|  | 9- to 12-year-old | 990 ± 265 | 868 ± 252 | 1103 ± 185 |
|  | 13- to 16-year-old | 942 ± 189 | 1169 ± 159 | 1097 ± 174 |
| PINV left response condition | 5- to 8-year-old | 1040 ± 161 | 1019 ± 193 | 1020 ± 195 |
|  | 9- to 12-year-old | 898 ± 210 | 957 ± 209 | 1029 ± 138 |
|  | 13- to 16-year-old | 912 ± 246 | 1054 ± 242 | 1094 ± 219 |

Table 3 Mean LRP amplitude values [µV] ± standard deviation of alpha-ERD for each CNV component. Significances (*p* < 0.5) are indicated for the respective values.

| Alpha-ERD LRP of CNV Component | Age Group | M1left/ M1right [µV] |
| --- | --- | --- |
| iCNV right response condition | 5- to 8-year-old | -12.73 ± 14  *t*(-3.14); *p* = .009 |
|  | 9- to 12-year-old | -9.79 ± 10.64  *t*(-3.8); *p* = .002 |
|  | 13- to 16-year-old | -5.83 ± 8.67  *t*(-3.77); *p* = .002 |
| lCNV right response condition | 5- to 8-year-old | -24.43 ± 27.97  *t*(-3.82); *p* = .003 |
|  | 9- to 12-year-old | -27.22 ± 33.05  *t*(-3.4); *p* = .004 |
|  | 13- to 16-year-old | -23.25 ± 20.22  *t*(-4.45); *p* < .001 |
| PINV right response condition | 5- to 8-year-old | -9.4 ± 12.19  *t*(-6.67); *p* = .02 |
|  | 9- to 12-year-old | -2.69 ± 9.75 |
|  | 13- to 16-year-old | -5.83 ± 8.67  *t*(-2.6); *p* = .02 |

___________________________________________________________________________

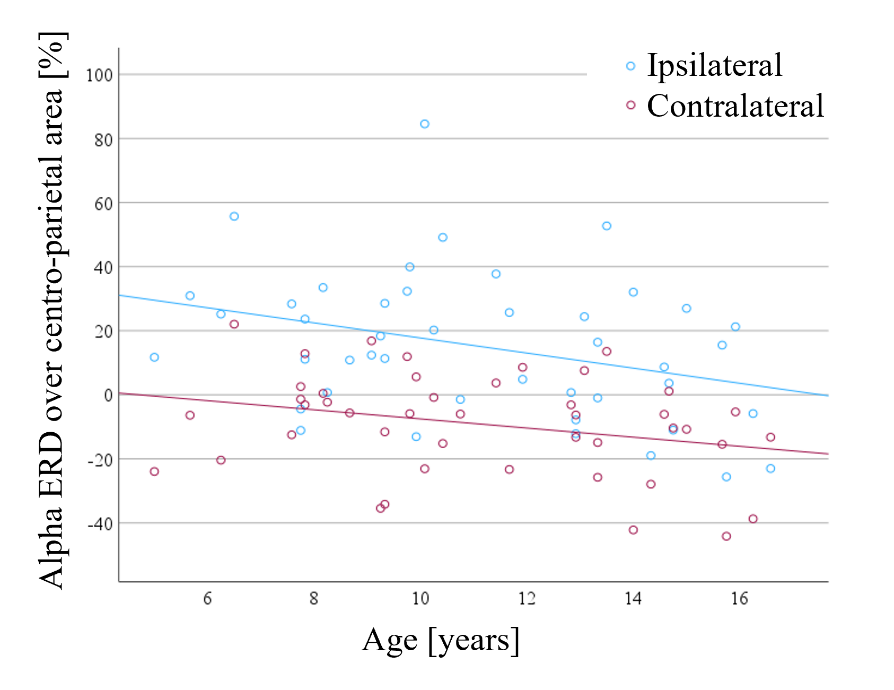

**Fig. 1.** Scatterplot of alpha power changes over centro-parietal scalp areas. Data points represent mean values of alpha ERD (%) for both response conditions and each subject, respectively for the ipsilateral (blue dots) and contralateral hemisphere (red dots). The associated colored lines show the fitting of a linear regression with an age-related decrease of ERS over ipsilateral motor areas and an age-related increase of ERD over contralateral motor areas.
___________________________________________________________________________
